# Tipping points in spatial ecosystems driven by correlated noise

**DOI:** 10.1101/2022.06.22.497206

**Authors:** Krishnendu Pal, Smita Deb, Partha Sharathi Dutta

**Affiliations:** Department of Mathematics, Indian Institute of Technology Ropar, Rupnagar 140 001, Punjab, India

## Abstract

Complex spatial systems can experience critical transitions or tippings on crossing a threshold value in their response to stochastic perturbations. Whilst previous studies have well-characterized the impact of white noise on tipping, the effect of correlated noise in spatial ecosystems remains largely unexplored. Here, we investigate the effect of both multiplicative and additive Ornstein-Uhlenbeck (OU) correlated noise on the occurrence of critical transitions in spatial ecosystems. We find that decreasing the noise correlation aggravates the chance of critical transitions - opposite to what is observed previously in temporal systems in the presence of OU-noise with fixed lag-1 correlation. Our results hold good and are supported by the analysis of three well-studied spatial ecological models of varying nonlinearity. We also compute spatial early warning indicators (e.g., spatial variance, spatial skewness, and spatial correlation) to determine their reliability in anticipating tipping points with variations in noise correlation. The indicators of critical transitions exhibit mixed success in forewarning the occurrence of a tipping point, as indicated by the distribution of Kendall’s rank correlation.

## I. INTRODUCTION

Due to unprecedented global environmental changes, the anticipation of critical transitions in spatial ecosystems is compelling. Critical transitions to a contrasting state have been identified in a plethora of complex spatial ecosystems including grasslands [1, 2], arid ecosystems [3], and vegetation colonization systems [3]. Research on ecosystems linking spatial pattern formation to catastrophic shifts unveils the importance of studying systems dynamics at a spatial scale [3–5]. For instance, characteristic spatial spotted vegetation pattern formation in various vegetative and grassland ecosystem models have signalled sudden transitions to a homogenous barren state [1, 4, 6, 7]. Investigating the spatial patterns of systems and the manifestation of such information provides early warnings of critical transitions [5] - ignoring which we may miss out on information essential to detect sudden transitions. Early warning indicators of critical transitions are designed to detect a dynamical phenomenon known as critical slowing down (CSD). At the onset of a tipping or bifurcation point, the system’s rate of return to the current equilibrium state upon a stochastic perturbations decreases as the dominant eigenvalue approaches zero, known as CSD [8]. CSD-based indicators, such as spatial variance, spatial skewness, and spatial autocorrelation at lag-1, are widely used to forewarn critical transitions in spatial systems [5, 9, 10] and exhibit an increasing trend prior to a transition.

Many studies have focused on resilience and sudden transitions in simple ecological models where environmental disturbances are modeled as white noise or noise with zero correlation [11, 12]. Whilst fluctuations depicted by white noise manifest ideal scenarios, in reality fluctuations are correlated [13–15]. Empirical evidence confirms that time series of various environmental variables often exhibit the presence of colored noise [16]. For improved prediction of catastrophic events, considering the paradigm of correlated noise is essential [17]. While white noise is characterized by frequencies having the same variance, correlated noise is characterized by non-stationary variance at different frequencies [16, 18]. Previous studies modeled extrinsic noise considering only the white spectrum to carry out a general theoretical study [19–21]. An essential factor that needs consideration is the attribute color of noise or modeling fluctuations in dynamical systems over the entire noise spectrum [22, 23]. In natural systems, perturbations are often not solely random but correlated with environmental variables with a finite correlation time [16, 24]. As the time scale of fluctuation exceeds the natural time scale of the dynamical system, the noise no longer remains independent and is unwise to be modeled by white noise. Such noises are correlated across time.

In the recent past, researchers have confirmed the influence of correlated noise in ecosystems [15, 16, 25], and marine events [23] such as heat storms and the presence of highly correlated noise in fish stocks. Lawton [26] found that populations perturbed with correlated noise stand a higher extinction risk than white noise perturbations. Vasseur and Yodzis [16] investigated the influence of noise color on many environmental variables. This scattered empirical evidence has guided the modeling of ecological and climate systems perturbed with correlated noise. Hitherto, little is known about the impacts of noise correlation on system dynamics envisioned at a spatial scale. Van der Bolt *et al*. [15] have shown that lag-1 autocorrelated noise increases the chance of tippings in temporal ecosystems. However, impacts of time-correlated noise in spatially extended ecological systems remain largely unknown owing to scanty literature and a lack of empirical evidence. Under this backdrop, we aim to study how environmental color noise affects critical transitions in spatial ecosystems.

To our knowledge, no studies have yet investigated the effects of correlated noise on the persistence of spatial ecosystems. Here, we study tippings in three well-studied spatial ecological models under the influence of correlated noise. The considered models are a vegetation-grazing model [27], a vegetation-turbidity model [28], and a lake-eutrophication model [29]. We incorporate the Ornstein-Uhlenbeck [30] color noise in the considered systems following a similar approach as used in non-equilibrium statistical mechanics. Within our framework, we aim to derive a general theory - of how noise correlation (weak → intermediate → strong) delays or precedes tippings in spatial ecological systems. We also address the impacts of noise correlation on the formation and width of the positive feedback/hysteresis loop in trying to recover species post-collapse by improving ecosystems conditions. Further, we calculate spatial EWSs for snapshots obtained from simulations of the three spatial ecological models for different noise realizations. We analyze the robustness of spatial early warning indicators in forecasting color noise-induced sudden transitions. Knowing the impact of noise correlation in spatially extended models will serve as a prelude to studying noise characterization in real spatial systems [31] aimed at evading or delaying tippings.

## II. MODELS AND METHODS

### A. Mathematical models

We adopt three well-studied ecological models which exhibit bistable dynamics via a saddle-node bifurcation to investigate the effect of OU-color noise in critical transitions. The first model in consideration is a population model in the presence of grazing pressure (see Eqn. (1)) [10] which has been used to study a variety of systems such as vegetation in a semiarid ecosystem [27], exploitation of fish communities [23], and spruce bud-worm dynamics [32]. Here, we consider a spatially extended version of this model. From now on, we shall refer to this model as the *vegetation-grazing* model. It describes the transition from a vegetated state to an over-exploited state as grazing pressure crosses a critical threshold. The model is expressed as:

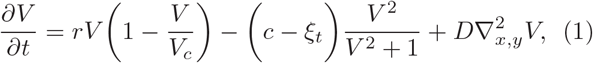

where *V* ≡ *V* (*x, y, t*) is the vegetation density and 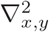 is the two-dimensional Laplacian operator working on *V*. *ξ*_*t*_ ≡ *ξ*(*x, y, t*) is the OU-color noise, multiplicative in nature here. Other model parameters are: the maximum growth rate *r*(= 10), the carrying capacity *V*_*c*_(= 10), and the maximum grazing rate *c*(= 15 − 30). The initial population *V*_*in*_ in all the cells are set in the same state and are sampled in the interval [10, 10.1].

We deploy a second model - the *vegetation-turbidity* model (Eqn. (2)), which describes the transition of the macrophytes-dominated clear water state of a shallow lake to the macrophytes-absent turbid state of water. Turbidity reduces vegetation density. This model describes the interactions between macrophyte coverage and turbidity of a shallow lake [9, 28]. The model is expressed as:

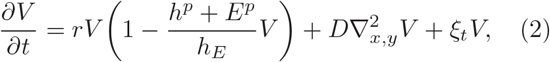

with 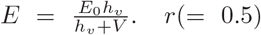 is the maximum growth rate, *h*(= 1.5) is the half saturation turbidity constant, *E*_0_(= 2 − 12) denotes the background turbidity, *h*_*v*_(= 0.2) is the half saturation vegetation cover, *p*(= 4) is the Hill coefficient. The initial vegetation *V*_*in*_ in each cell is sampled randomly in the interval [0.5, 0.6].

The third model we choose describes the nutrient dynamics of a eutrophic lake [29]. When the nutrient input rate is low, the lake remains oligotrophic as the nutrients are lost due to sedimentation. However, when nutrient loading increases, nutrients recycle from the sedimentation back to the water by high recycling due to lower oxygen levels, and the lake suddenly becomes eutrophic [9]. Now onwards we refer to it as the *lake-eutrophication* model (Eqn. (3)). The model is expressed as shown below:

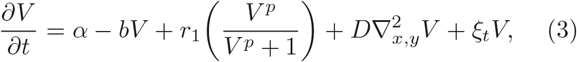

where we choose the nutrient loading rate *α*(= 0.1 − 0.9), the nutrient loss rate *b*(= 1), the maximum recycling rate *r*_1_(= 1), and the Hill coefficient *p*(= 8). The initial vegetation in each cell is sampled randomly in the interval [0.4, 0.5].

In all the aforementioned three models, *D*(= 0.001) is the diffusion/dispersion rate, spatial resolution *dx* = *dy* = 0.1, time increment *dt* = 10^−3^, spatial grid size (*n*_*x*_ × *n*_*y*_ = 50 × 50) and *ξ*(*x, y, t*) is the OU-color noise whose properties will be described in the next subsection.

### B. Ornstein-Ulhenbeck noise

For generating color noise we consider the Ornstein-Uhlenbeck [33, 34] process. The OU-colored noise or OU-noise *ξ*(*t*) is calculated from the equation:

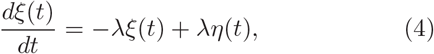

where *ξ*(*t*) is the OU-noise and *η*(*t*) is Gaussian white noise source with ⟨*η*(*t*) ⟩ = 0, and ⟨*η*(*t*)*η*(*t*′)⟩ = 2*D*_0_*δ*(*t* − *t*′). The solution of the Eqn. (4) satisfies:

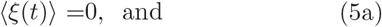

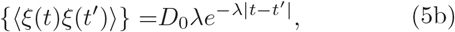

where Eqn. (5b) is the correlation of the OU-noise, *λ*^−1^ is called the correlation time, and *D*_0_ is the noise amplitude. The {‥} denotes the averaging over the distribution of initial *ξ*_0_ values, which is given by:

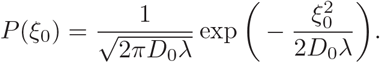

The secondary averaging {‥} is essential for stationary correlation given in Eqn. (5b). The Eqn. (5b) often alternatively written as:

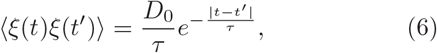

where *τ* = 1*/λ* is called correlation time. In our spatially extended system as the noise in individual grid is only temporally correlated and spatially uncorrelated, the better notation of noise correlation than Eqn. (6) would be as follows:

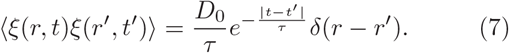

From now on, while discussing about the noise correlation we shall refer to the Eqn. (7). Thus the OU–noise has a nonzero correlation time *τ*, meaning that values of the noise at different times are not independent random variables. Further, from Eqn. (6) it is clear that in the limit *τ* → 0, the autocorrelation approaches 2*D*_0_*δ*(*t* − *t*′) and the noise *ξ*(*t*) tends to the Gaussian white noise *η*(*t*).

### C. Color noise generation

We begin with the standardization of the OU-noise generation. The correlation of the noise is given by Eqn. (7). The Integral method of generating OU-noise is discussed in detail in Section II D. In Fig. 1 we show the normalized correlation for five cases up to |*t* − *t*′|= 5, i.e., up to 5000 lag as subsequent noises are separated by *dt* = 10^−3^ time unit for all the three models. Each of the simulated trajectories is an average of 125 independent simulations of Eqn. (4), and each trajectory is a sequence of 10^4^ data points. The solid and dotted lines indicate the numerically simulated normalized correlation, and the normalized analytical correlation derived from Eqn. (7). We have plotted the curves with increasing correlation time (τ) starting from τ = 0.1 to τ = 5.0. One can observe from Fig. 1 that as the correlation time increases, the correlation or color of the noise increases. Fig. 1 confirms that the noise generated by the above method accurately generates OU-noise. However, accuracy may further be improved with consideration of more data points. For our purpose, considering 104 data points is sufficient.

**FIG. 1.**
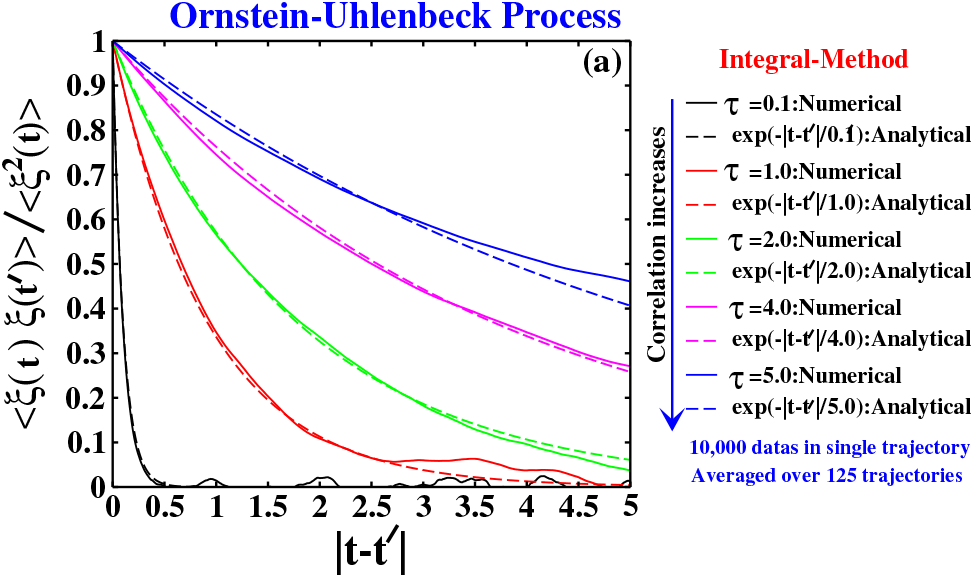
Standardization of the OU-noise: The analytical expression of OU-noise correlation plotted against the numerically obtained normalized correlation using the integral method for five cases ranging from *τ* = 0.1 to 5. As depicted with an increase in the correlation time *τ*, the correlation or color of the noise increases. In the limit *τ* → 0, the noise resembles the Gaussian white noise *η*(*t*).

### D. Numerical simulation: Spatial models

To simulate the models (Eqns. (1), (2), and (3)), we employ Euler’s FTCS (Forward in Time and Central in Space) method using FORTRAN 95 programming language. We first construct the spatial grid of dimension (*n*_*x*_ × *n*_*y*_ = 50 × 50) with spatial resolution *dx* = *dy* = 0.1. We use periodic boundary condition for all model simulations. We provide the initial values of the state variable *V*_*in*_ with random perturbation (*q*) in each grid, *q* ∈ [0, 0.1] to break the square symmetry. We then perform integration in each grid cell with integration time step *dt* = 10^−3^ incorporating OU-noise as:

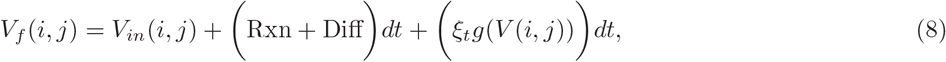

with 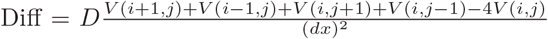, Rxn = *f* (*V* (*i, j*)), *V*_*f*_ is the final value upon integration of *V* (*i, j*) after each iteration with previous value of *V* (*i, j*) denoted by *V*_*in*_, and *g*(*V*) is the multiplication factor of the state variable *V*. *ξ*_*t*_ is the individual OU-noise in each grid. We run each model simulation up to time *T*_*s*_, where the steady-state for the concurrent bifurcation parameter is already achieved apriori in the deterministic (noiseless) model. Furthermore, *T*_*s*_ has been chosen so that for all bifurcation parameters, a deterministic system gets enough time to go to a steady-state. We then use sticky initial conditions, i.e., the final spatial distribution is the initial condition for the next bifurcation parameter. This way, we run the simulations until done for the final value of a bifurcation parameter. We obtain the snapshots for each final spatial distribution of the state variable at each value of the bifurcation parameter.

For the OU-noise generation, we apply a widely used integral method [33]. Integrating equation (4) one obtains:

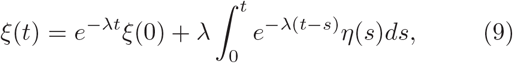

and

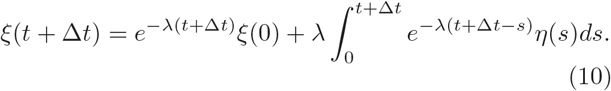

Subsequently,

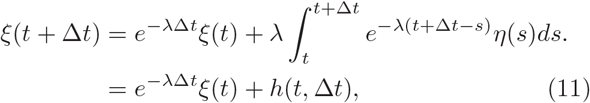

where the function *h*(*t*, Δ*t*) is Gaussian with zero mean as *η*(*t*) is Gaussian with zero mean. Therefore, all its properties are determined by its second moment:

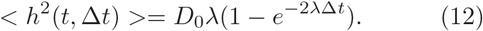

To start the simulation an initial value for *ξ* is required and that is obtained as follows: (i) First, we generate two random numbers *r*_1_[0,1] and *r*_2_[0,1]. (ii) Then the initial value of *ξ*(J*t*) is obtained from a Gaussian random number 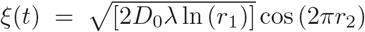. This is exactly the Box-Muller Algorithm [35] for generating Gaussian random number. After that the exponentially correlated noise is obtained as follows: (iii) We call another two random numbers *r*_3_[0, 1] and *r*_4_[0, 1]. (iv) Then we set 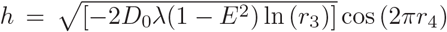, where *E* = exp(−*λ*Δ*t*). (v) Now the OU-noise is obtained by *ξ*_*t*+*dt*_ = *ξ*(*t*)*E* + *h*. (vi) Finally, this process loops back to step (iii) and continues as long as required.

### E. Spatial early warning signals (S-EWSs)

Next, we calculate the S-EWSs [5] such as spatial standard deviation, spatial skewness, and spatial auto-correlation using the spatial early warning signal package (www.early-warning-signals.org). Close to the bifurcation point, the system’s equilibrium state experiences more substantial fluctuations [36] and computing S-EWSs in such instances and investigating their efficacy is essential. Spatial variance (*σ*^2^) is the second moment around the spatial mean of the state variable *V* and is defined as:

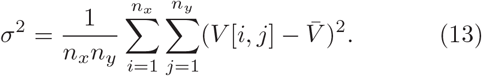

The square root of the variance: *σ* is called the spatial standard deviation [5].

As the system approaches a tipping point, the fluctuations around the mean can also become increasingly asymmetric. This is because fluctuations in the forward direction to the alternative stable state take longer to return to equilibrium than those in the backward direction [10]. This spatial asymmetry is measured by spatial skewness (*γ*), which is the third central moment scaled by the standard deviation [5], and defined as:

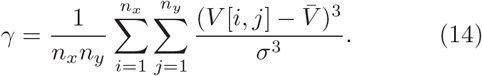

When a system approaches a tipping point, the spatial coupling force between adjacent units becomes dominant due to increased recovery time to local equilibrium. These neighboring units become increasingly correlated [9]. The increasing spatial coherence is quantified by the spatial correlation function *C*_2_(*r*), or Moran’s I [5], between ecological states *V* [*i, j*] and *V* [*m, n*] separated by a distance *r*. The spatial correlation function *C*_2_(*r*) is defined as below:

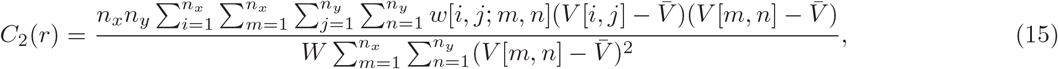

where 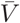 is the spatial mean of the state variable 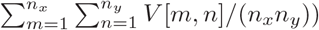. *w*[*i, j*; *m, n*] takes value 1 if positions [*i, j*] and [*m, n*] are separated by a distance *r*, and assumes the value 0 otherwise. *W* is the total number of units separated by distance *r. C*_2_(*r*) is the correlation in the two-dimensional plane. When the system approaches a tipping point, critical slowing down becomes more pronounced and the spatial correlation function increases at all distances [9].

## III. RESULTS

### A. Critical transitions in the presence of OU-noise

We study the dynamics of the three spatial models (Eqns. (1), (2), and (3)) perturbed with multiplicative OU-noise and observe the variation in tipping for change in the correlation time *τ* of the OU noise. The first model in consideration is the vegetation-grazing model (Eqn. (1)). In Fig. 2(a), we have plotted the mean vegetation density 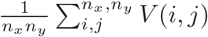 against increasing grazing rate *c* for different correlation time *τ*. The black-dashed line shows the mean density in absence of any noise, i.e., the deterministic mean density. The mean density has been plotted for decreasing *τ* values ranging from 10.0 to 0.01. We observe advanced critical transition with decreasing noise correlation, i.e., the threshold value of the bifurcation parameter (*c*) at which the tipping occurs is comparatively smaller for lower *τ* values. The threshold value of the driver at which critical transition occurs for the two extreme *τ* values; such as for *τ* = 10.0 and *τ* = 0.01 is significantly altered in the presence of color noise.

**FIG. 2.**
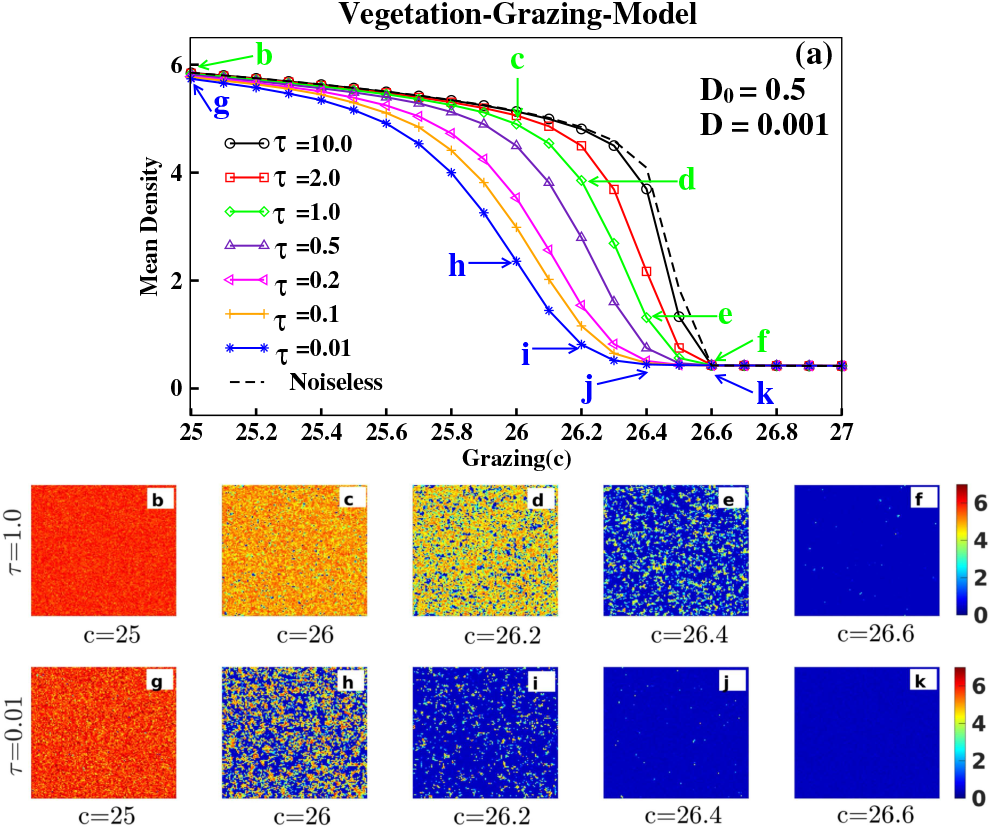
Critical transition in the vegetation-grazing model (Eqn. (1)): (a) Spatial mean vegetation density plotted against gradual change in the grazing rate *c*. The system undergoes delayed critical transitions for higher *τ* values (correlation time) from the higher stable (density) state to the lower stable state compared to lower *τ* values. The black dashed trajectory represents the density of vegetation in the deterministic system. (b)-(k) Spatial vegetative patterns observed at different values of the grazing rate *c* as marked in (*a*). The color bar indicates the density of vegetation. All parameter values in the above, as well as other figures unless mentioned, take the same values, and the constants *D*_0_, *D* have the same meaning as described in subsection II A.

For a more clear understanding of the aggravated critical transition for reduced *τ* values, we visualize sequences of spatial patterns at two *τ* values (*τ* = 1.0 and *τ* = 0.01). Figs. 2(b)-2(f) and Figs. 2(g)-2(k) represent the sequence of 5 snapshots for grazing rates *c* = 25, 26, 26.2, 26.4 and 26.6 for *τ* = 1.0 and *τ* = 0.01, respectively. The respective snapshots that we have considered are marked in the mean density plot (Fig. 2(a)). Comparing Figs. 2(b)-2(f) and Figs. 2(g)-2(k) one can observe that the threshold for extinction of the vegetation population is *c* = 26.4 for *τ* = 0.01 (see Fig. 2(j)), while there is significant amount of vegetation remaining at this *c* value for *τ* = 1.0 (see Fig. 2(e)). At *c* = 26.6 the system shows barren state for both the *τ* values (Figs. 2(f) and 2(k)). Thus it is evident from both the mean density plots and the spatial pattern visualizations, with decreasing color noise correlation the critical transition occurs earlier in the spatial vegetation population.

Next we study the vegetation-turbidity model (Eqn. (2)) where increasing background turbidity (*E*_0_) serves as the bifurcation parameter which turns the water of a shallow lake from the clear state to a turbid state. Fig. 3(a) shows the mean vegetation density plotted as a function of increasing *E*_0_ for different noise correlation ranging from *τ* = 100.0 to *τ* = 2.0. In Figs. 3(b)-3(f) and Figs. 3(g)-3(k) we have plotted the spatial distribution of the vegetation population for background turbidity *E*_0_ = 5.1, 5.3, 5.4, 5.5 and 5.6 for *τ* = 100.0 and *τ* = 2.0, respectively. Spatial distribution in Fig. 3(j) shows that for *τ* = 2.0 the vegetation is in the verge of extinction, while Fig. 3(e) shows that for *τ* = 100.0 sufficient vegetation density still persists at *E*_0_ = 5.5.

**FIG. 3.**
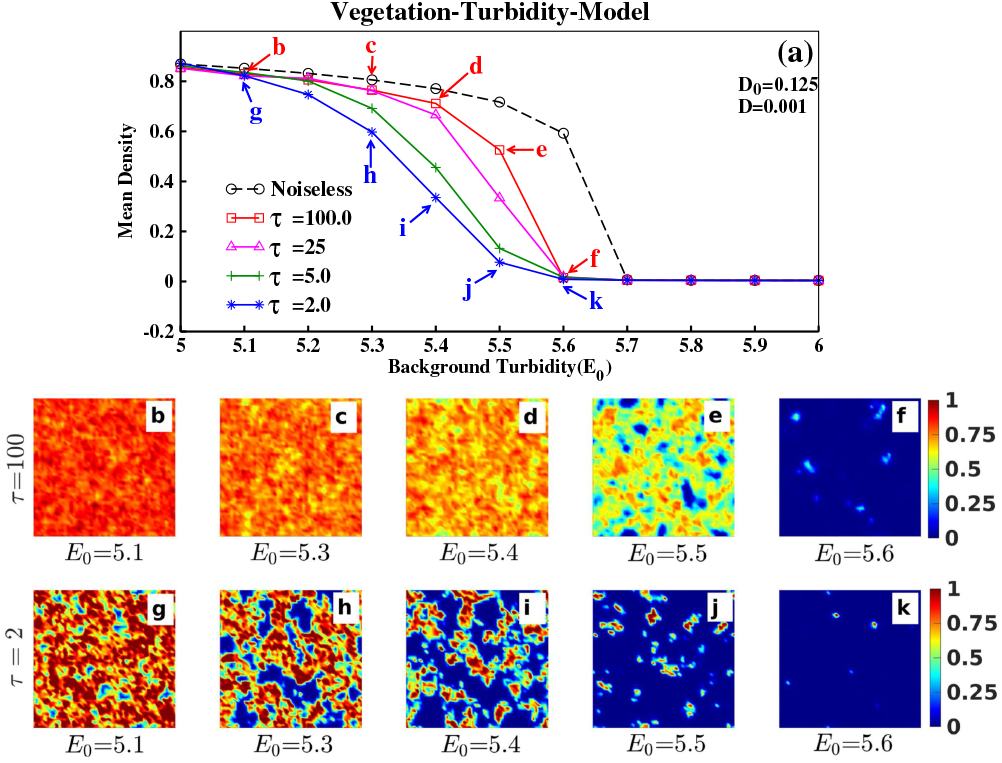
Critical transition in the vegetation-turbidity model (Eqn. (2)): (a) Spatial mean vegetation density plotted against gradual change in the background turbidity *E*_0_. The system undergoes comparatively delayed sudden transitions from the higher stable (density) state to the lower stable state for higher correlation time *τ*. The black dashed trajectory represent the density of vegetation in the deterministic system. (b)-(k) Spatial vegetative patterns observed at different *E*_0_ values as marked in (*a*). Color bar indicates the density of vegetation.

We analyse the third model, i.e., the lake-eutrophication model (Eqn. (3)). As the control parameter - nutrient loading rate (*α*) increases, the lake transforms from a clear lake state to an eutrophic lake state with increasing vegetation density in the lake. Fig. 4(a) represents the mean vegetation density with the increasing nutrient loading rate in presence of OU-noise with correlation time ranging from *τ* = 20.0 to *τ* = 1.0.

**FIG. 4.**
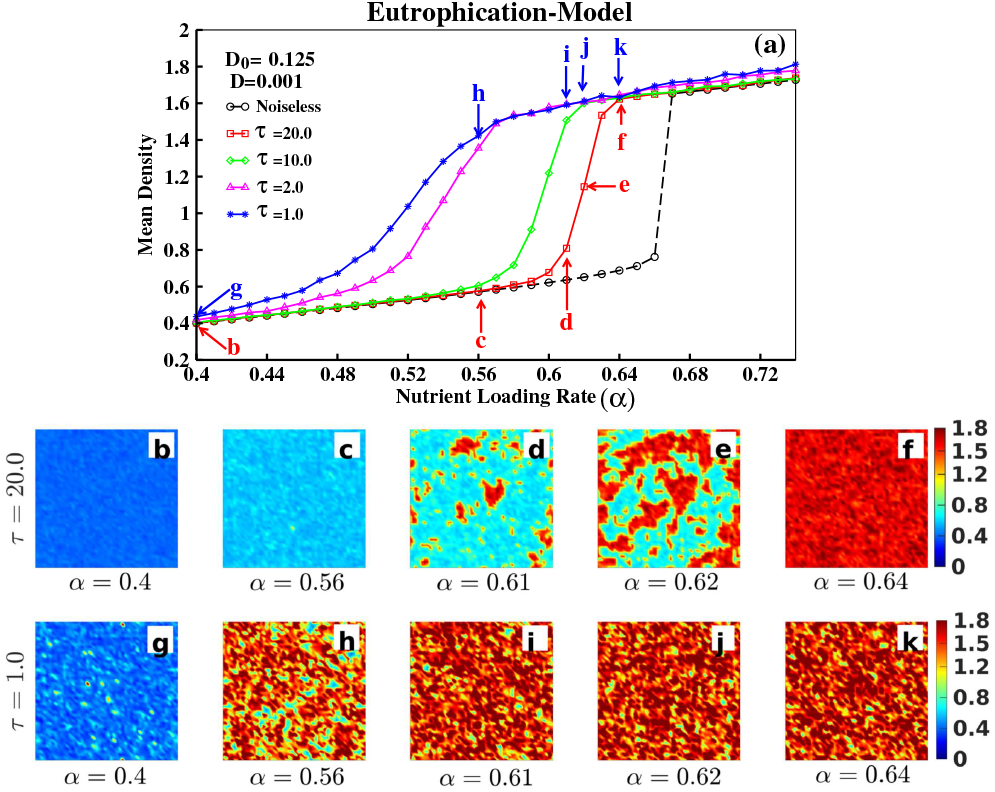
Critical transition in the lake-eutrophication model (Eqn. (3)): (a) Spatial mean nutrient concentration plotted against gradual change in the nutrient loading rate *α*. The system undergoes delayed sudden transitions from the lower stable (density) state to the higher stable state for higher correlation time *τ* than for lower *τ* values. The black dashed trajectory represents the density of nutrients in the deterministic system. (b)-(k) spatial vegetative patterns observed at different *α* values as marked in (*a*). The color bar indicates the nutrient concentration in the lake.

The spatial distribution of the vegetation density for nutrient loading rates *α* = 0.4, 0.56, 0.61, 0.62 and 0.64 for two cases, such as *τ* = 20.0 and *τ* = 1.0, are plotted in Figs. 4(b)-4(f) and Figs. 4(g)-4(k), respectively. From left to right spatial snapshots the lake becomes increasingly eutrophic as displayed in Fig. 4. For *α* = 0.56 and *τ* = 1.0, Fig. 4(h) indicates a eutrophic lake. While at *α* = 0.56 and *τ* = 20.0, Fig. 4(c) indicates that the lake is yet to become fully eutrophic and still persists. Thus our results are robust across models of varying nonlinearity and indicate that increase in noise correlation time *τ* can delay a critical transition.

### B. Hysteresis loop

In this section, we elucidate on the presence of a hysteresis loop, if any. As a control parameter is gradually varied, all the models pass through a saddle-node bifurcation resulting in a transition to an alternate state. Striving to recover system post-collapse, we observe the presence of alternate attractors, the trajectory which the system follows on improving conditions. Thus the point of collapse and recovery are not same and this results in the formation of a hysteresis loop. Moreover, the formation of a hysteresis loop also confirms that the observed transitions are first-order transitions [37]. In Figs. 5(a)-5(c) we show the occurrence of critical transition in spatial mean density in the forward direction (in solid) for deterioration in environmental conditions for all the considered spatial models, respectively. By improving conditions through gradually varying the control parameter in the backward direction we show the point of recovery for all the models. In Fig. 5(a), we plot the spatial mean density for linear increase (forward) and decrease (backward) in the control parameter (*c*) and observe the formation of hysteresis loops for *τ* = 1.0 and *τ* = 0.01, respectively. As previously depicted in Fig. 2, as *τ* decreases the critical transition occurs early in the forward direction. Similarly, the system experiences an early recovery for lower value of the correlation time. Thus, increasing the value of *τ* the width of the hysteresis loop increases.

**FIG. 5.**
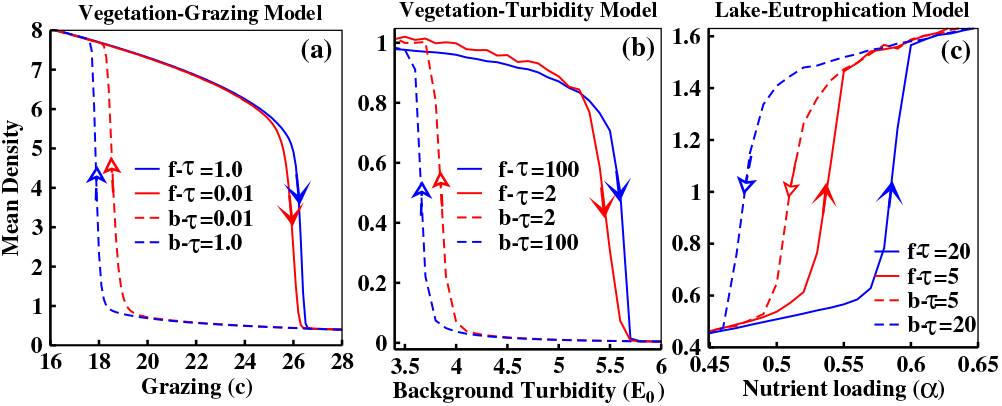
Catastrophic collapse and recovery in the ecosystem models: On increasing the control parameter/ecosystem driver in the forward direction, a system undergoes a catastrophic shift to an alternative state. While decreasing the driver in the opposite direction, striving to recover the system post-tipping, the system exhibits an abrupt shift in density beyond a critical value of the control parameter, leading to the formation of a hysteresis loop. The width of the hysteresis loop increases with the correlation time *τ*. The red (blue) trajectories in each sub-figure correspond to low (high) *τ* values.

In Fig. 5(b) we have shown similar plots for *τ* = 100 and *τ* = 2 and in Fig. 5(c) the similar cases have been shown for *τ* = 20 and *τ* = 5. From Fig. 5, it is evident that with increasing noise correlation, the width of the hysteresis loop increases, indicating that with increasing noise correlation the transitions will be delayed.

### C. Transitions driven by additive coloured noise

For additive coloured noise, the Eqn. (8) changes to the following form:

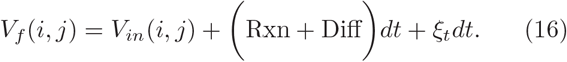

We have found similar results as in cases of multiplicative noise for all the models. For an example, in Fig. 6 we show the results for the lake-eutrophic model (Eqn. (3)): We observe the mean density plots for each noise realization in Fig. 6 and are conclusive of similar trends of delayed critical transition in the system on perturbing with additive OU-noise with higher correlation time *τ*.

**FIG. 6.**
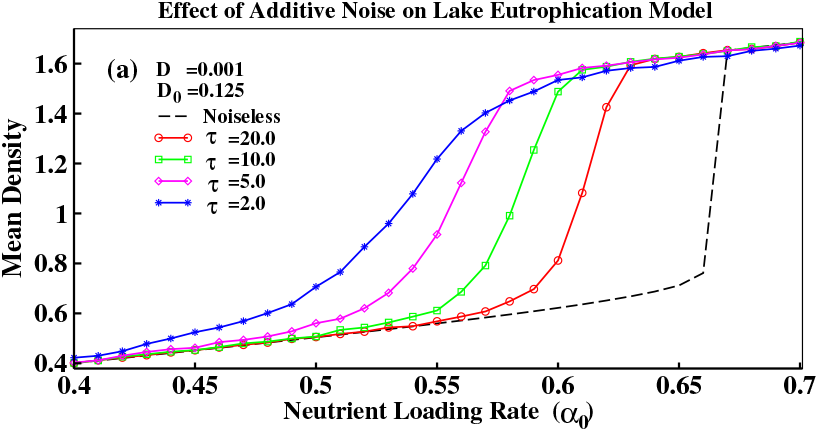
Critical transition under the influence of additive coloured noise: Spatial mean density plotted for gradual change in nutrient loading rate for different *τ* values. System exhibit dynamics are similar to that of Fig. 4 where the system is perturbed with multiplicative noise. The black dashed trajectory represents the density of vegetation in the deterministic system.

### D. Spatial early warning signals analysis

In this subsection we study the effect of noise correlation on the efficacy of spatial early warning signals in forecasting a sudden transition. We have calculated the three leading S-EWSs such as spatial standard deviation, spatial skewness and spatial lag-1 autocorrelation using the equations shown in the subsection II D. In the top panel of Figs. 7(a)-7(c) the spatial standard deviations are shown for all the three models, respectively. A rising trend in spatial standard deviations is observed for the sequence of pre-transition snapshots from all the three models for different *τ* values. With decreasing noise correlation as the tipping occurs early the increment in the standard deviation is also gradually shifting towards left indicating an early tipping with decreasing noise correlation. In the middle panel of Figs. 7(d)-7(f) the trends in spatial skewness are plotted. The peaks here also shifts towards left indicating early tipping with decreasing noise correlation. In the bottom panel, Figs. 7(g)-7(i) the spatial Lag-1 auto-correlations have been plotted which also shows similar trends. Thus from the S-EWSs analysis in Fig. 7, it is confirmed that the delay in critical transitions due to decreasing noise correlation is correctly being captured. The strength of the indicators are evaluated by calculating the Kendall’s-*τ* rank correlation coefficient for each sequence of snapshot corresponding to the three models for all the noise realizations are shown in Table I. The S-EWSs analysis done here corroborates with results that we have shown above.

**FIG. 7.**
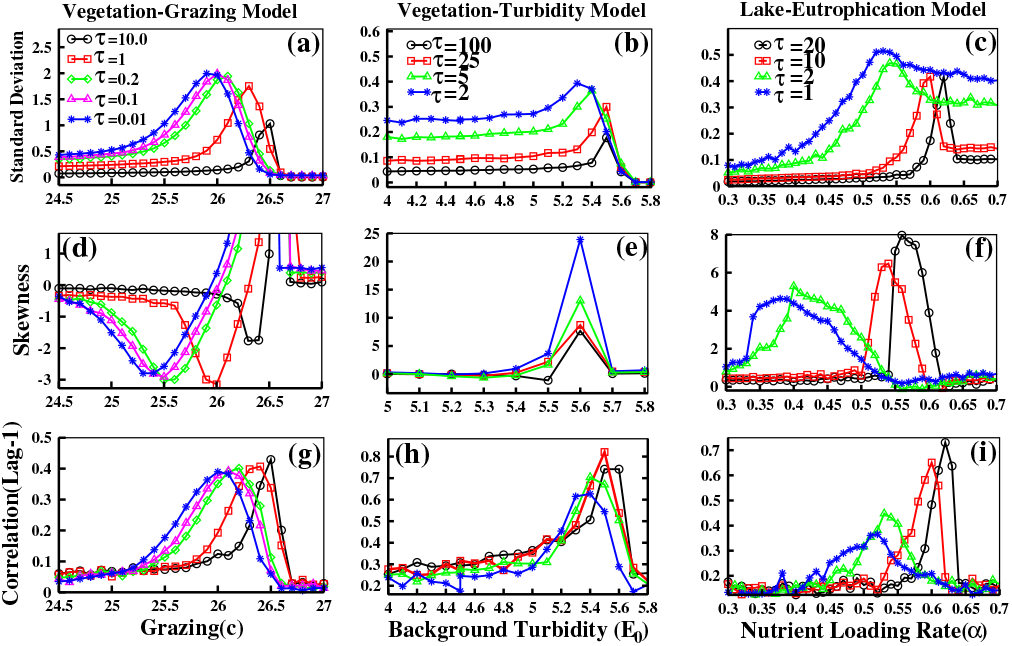
Trends in generic spatial indicators along change in degradation parameter across all the considered spatial models: (a)-(c) Spatial standard deviation, (d)-(f) Spatial skewness, and (g)-(i) spatial correlation at lag-1 obtained from snapshots along the gradients of the control parameter for different *τ* values for each model. The indicators are calculated using 80, 60, and 61 snapshots for the considered models, respectively. Spatial standard deviation and correlation at lag-1 exhibit a rising trend on approaching tipping for all *τ* values irrespective of the model chosen. On the other hand, mixed trends are observed for spatial skewness across models and *τ* values.

**TABLE I.** Kendall’s-*τ* statistics for spatial early warning indicators: The table below presents Kendall’s-*τ* correlation coefficient for spatial indicators viz spatial standard deviation, spatial skewness, and spatial correlation, respectively corresponding to spatial snapshots generated from the three considered models perturbed with correlated noise with different correlation time *τ*.

| Models and parameters | Spatial variance | Spatial skewness | Spatial correlation |
| --- | --- | --- | --- |
| Vegetation-Grazing |  |  |  |
| $\tau = 0.01$ | 0.992 | -0.826 | 0.75 |
| $\tau = 0.1$ | 0.992 | -0.827 | 0.7 |
| $\tau = 0.2$ | 0.992 | -0.84 | 0.66 |
| $\tau = 1$ | 0.991 | -0.814 | 0.673 |
| $\tau = 10$ | 0.997 | -0.69 | 0.74 |
| Vegetation-Turbidity |  |  |  |
| $\tau = 2$ | 0.64 | -0.33 | 0.54 |
| $\tau = 5$ | 0.794 | -0.413 | 0.70 |
| $\tau = 25$ | 0.75 | -0.625 | 0.737 |
| $\tau = 100$ | 0.80 | -0.238 | 0.70 |
| Lake-Eutrophication |  |  |  |
| $\tau = 1$ | 0.97 | -0.377 | 0.784 |
| $\tau = 2$ | 0.957 | -0.09 | 0.706 |
| $\tau = 10$ | 0.99 | -0.60 | 0.33 |
| $\tau = 20$ | 0.974 | -0.33 | 0.203 |

### E. Significance testing of the spatial early warning signals

The strength of S-EWSs in forecasting an impending transition can be quantified by calculating Kendall’s-*τ* rank correlation coefficient for the indicators for a large number of surrogate sequences of snapshots. Such significance testing helps identify false signals (Type-I and Type-II errors) and assure whether trends in indicators are random due to chance. Here, we specify a null hypothesis to compare with trend estimates of the original data and reject the null hypothesis if *p < α* (if the probability (*p*) of obtaining a value from the experiment is lower than the significance level (*α*), then we reject the hypothesis that such trends obtained - are not comparable to the trends for the original data and occur due to chance). We test the null hypothesis by generating 100 surrogate spatial snapshots for each realization of noise correlation for all the considered models. We observe the distribution of Kendall’s-*τ* values (ranging over -1 to 1) for each realization to comment on the occurrence of false signals of an upcoming abrupt transition. Box plots in Figs. 8(a)-8(c) indicate that standard deviation is a robust indicator and can reliably predict tipping in spatial systems when perturbed with OU-noise. The strength of the indicator does not alter with varying noise correlation *τ*. On the other hand, spatial lag-1 correlation exhibit stronger trends for all *τ* values in the vegetation-grazing model and moderate strength in the vegetation-turbidity model. For the lake-eutrophication model, we observe stronger trends for the same at *τ* = 1, 2 (see Fig. 8(h)). However, on increasing *τ* further, we observe weaker signals as indicated by the Kendall’s-*τ* distribution in Fig. 8(i). Contrarily, spatial skewness trends are weaker for all *τ* values, and we obtain no signals in some instances (see Figs. 8(d)-8(f)). All the trends obtained in Fig. 8 are significant with *p <* 10^−5^.

**FIG. 8.**
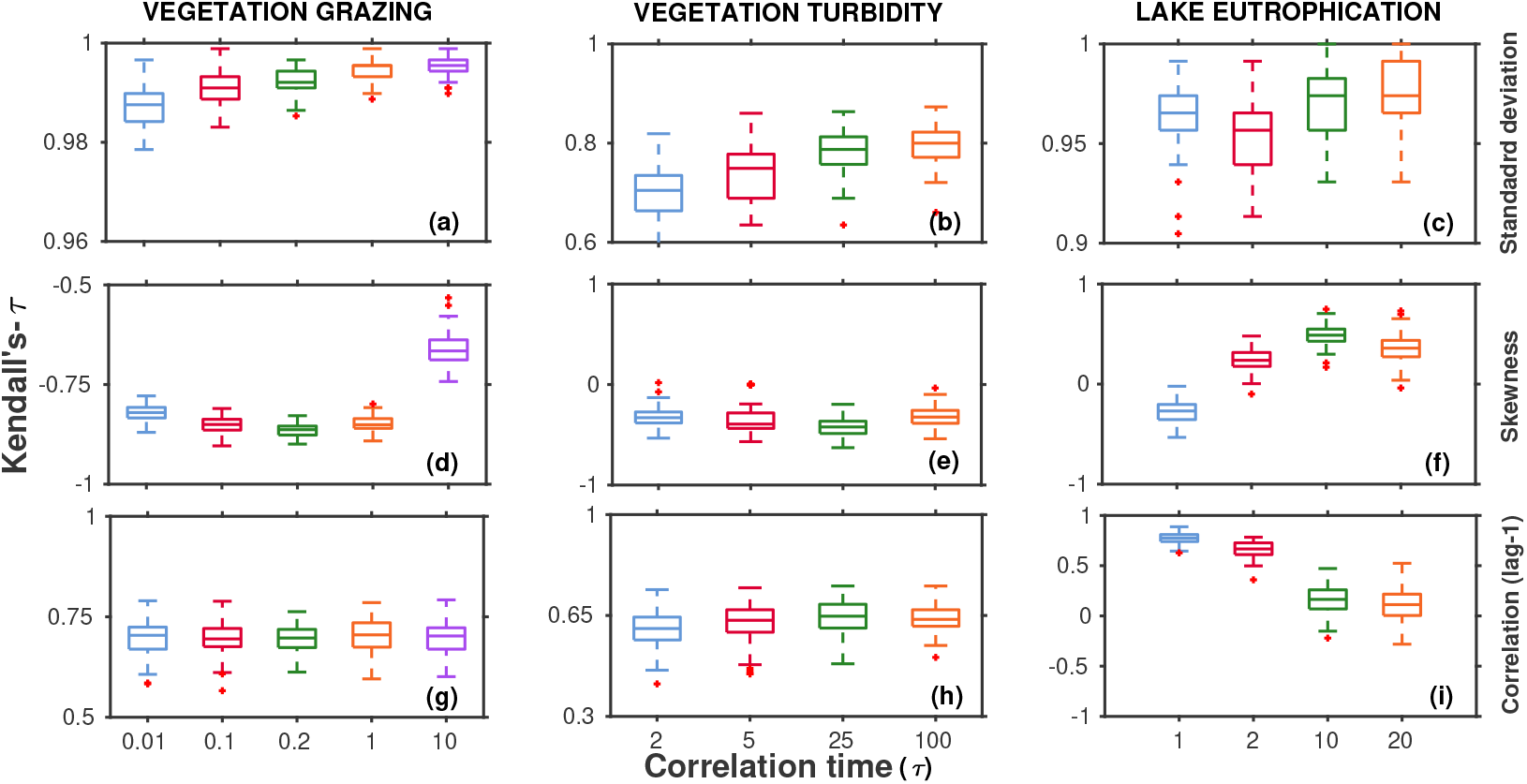
Kendall’s-*τ* test statistics for spatial early warning signals: Test statistics indicate the strength of trends in (a)-(c) spatial standard deviation, (d)-(f) skewness, and (g)-(i) correlation at lag-1, respectively. Box plots illustrate the distribution of Kendall’s-*τ* values obtained across 100 surrogate spatial snapshots corresponding to each correlation time *τ*. Standard deviation is the most robust indicators to detect sudden transitions for different correlated noise across different ecological models. Trends statistics for all the indicators are significant (*p <* 10^−5^).

## IV. CONCLUSIONS AND DISCUSSION

To rightfully anticipate tipping points in complex systems, bridging the gap between the dynamics of real-world spatial systems and that of simple models is essential. While this is being done, considering correlated noise is one of the few primary facets for mimicking dynamics of natural systems [16, 17]. Ecological systems may be affected by correlated noise where species dynamics are directly modulated, leading to random interaction changes within species. Although the later has an indirect impact on the dynamics, the effect of correlated noise on critical transitions of spatial systems is principal though less explored thus far. Here we have analyzed how critical transitions and related S-EWSs are influenced by exponentially correlated color noise generated from the OU process [33] on three spatial ecological models of different origins. We present a more general study considering the noise that is spatially uncorrelated and exponentially correlated in time, quite likely for environmental disturbances that may be correlated over varying delay times. Contrary to previous studies, which show lag-1 auto-correlated noise toward the red end of the noise spectra advance critical transitions [15, 38], our findings (not limited to only lag-1 auto-correlated noise) of advanced critical transitions at low *τ* values shall motivate researchers to cultivate further to comment on the generality of the statement. However, the threshold *τ* value, which leads to tippings, may differ across systems, and it depends on the timescale of the system. Clearly, from our result in Fig. 5, at a higher *τ* value, the width of the hysteresis loop increases, which implies that by improving the environmental conditions post tipping, recovery in systems is comparatively delayed for high *τ* values. Our results are qualitatively similar and hold good for all the models perturbed with both additive and multiplicative noise.

Further, our studies reveal that persistence of memory in fluctuation influences the occurrence of a critical transition in all the three spatial models of varying nonlinearities - a step ahead towards a general conclusion. However, robust indicators that forewarn such transitions are lacking, and results confirm that generic S-EWSs have had mixed success and their reliability reduced further with more correlated noise [38]. However, despite the challenges, recent work has sought to improve early warning signals using deep learning techniques [39]. While such methods are propitious, it requires further fine-tuning before such models serve as universal indicators of critical transitions. Owing to these concerns, we lay the foundation of a more in-depth study of such transitions in complex higher-dimensional spatial systems and provide robust indicators of the same. We can infer from our results in Table. I and Fig. 8 that S-EWSs, standard deviation, and correlation at lag-1 constitute robust indicators of critical transitions irrespective of the noise color in the system across models. However, spatial skewness exhibit mixed trends and weak signal strength.

The complexity of the diffusion equations limits the ability to provide explicit analytical expressions to support our numerical results. However, our noise implementation is standardized using integral and differential methods of generating OU-noise[33]. Further, the lack of empirical data availability limits the generality of our results based on model simulations, while real scenarios may manifest more complex dynamics. Nonetheless, our study implicates that increasing the noise correlation in a degraded system can delay the sudden transition. An interesting future work may include extending the study to a larger class of spatial systems and further investigating the interplay of diffusion rate and noise intensity in tippings. It may be noted that bistability is observed in patchy vegetative systems [3] where self-organized patterns signal the onset of a critical transition alongside trends in spectral EWSs. While this work focuses on pretransition non-homogeneous spatial patterns, a promising direction includes studying the impact of color noise in systems with self-organized patterns. This will be instrumental in concluding the effects of color noise in spatial ecosystems across diverse fields.

## ACKNOWLEDGMENTS

K.P. acknowledges IIT Ropar for postdoctoral fellowship (F.No.: 01-DR/2019/IITRPR/4952). S.D. acknowledges the Ministry of Education (MoE), Govt. of India for Prime Minister’s Research Fellowship (PMRF). P.S.D. acknowledges financial support from the Science & Engineering Research Board (SERB), Govt. of India [Grant No.: CRG/2019/002402].

